# Near-infrared optoacoustic modulation of the blood-brain barrier permeability using size-tuned hyperbranched gold nanoconstructs

**DOI:** 10.64898/2026.08.30.748173

**Authors:** Yu Wu, Yu Ge, Xitao Li, Hongxin Sun, Yejun Zhang, Chunyan Li, Guangcun Chen, Jiang Jiang

**Affiliations:** School of Nano-Tech and Nano-Bionics, University of Science and Technology of China, Hefei 230026, China; CAS Key Laboratory of Nano-Bio Interface, Suzhou Key Laboratory of Functional Molecular Imaging Technology, Division of Nanobiomedicine and i-Lab, Suzhou Institute of Nano-Tech and Nano-Bionics, Chinese Academy of Sciences, Suzhou 215123, China; College of Sciences, Shanghai University, Shanghai 200444, China

**Keywords:** hyperbranched gold, optoacoustic, femtosecond laser, blood-brain barrier

## Abstract

The blood–brain barrier (BBB) constitutes a major bottleneck for the systemic delivery of most therapeutic agents to the central nervous system. Here, we report near-infrared reversible optoacoustic modulation of the BBB permeability (NIR-ROAMBBB), leveraging endothelial tight junction targeting hyperbranched gold nanoconstructs (HBGNCs) to amplify localized optoacoustic transduction under femtosecond laser excitation. We first synthesized HBGNCs with tunable particle sizes (62–150 nm) and consistent branch morphologies via a seed-mediated growth approach, and uncovered a non-monotonic relationship between particle dimension and optoacoustic output, where the 62 nm HBGNCs generated nearly twofold stronger optoacoustic signal than gold nanorods and gold nanostars under matched excitations. Conjugation with BV11 antibodies against junctional adhesion molecule A increased HBGNC endothelial association and cerebral accumulation, enabling focal and fluence-dependent transient BBB opening (3-6 h) under 800 nm femtosecond pulsed laser excitation, as validated by in vitro trans-endothelial electrical resistance measurements, ex vivo Evans blue extravasation staining, and in vivo NIR imaging. Featuring deep tissue penetration of NIR light, robust optoacoustic conversion of HBGNCs, and negligible femtosecond laser-induced photothermal damage, this non-invasive strategy enables precise focal modulation of BBB permeability and potential drug delivery.

## Introduction

The blood-brain barrier (BBB) is a specialized neurovascular interface that preserves central nervous system homeostasis by tightly regulating molecular exchange between the circulation and neural tissue.^1–3^ However, this protective function also restricts brain exposure to many systemically administered drugs.^4, 5^ Spatially precise, temporally controlled, and reversible modulation of BBB permeability is therefore important for delivering therapeutics to treat central nervous system (CNS) diseases. Out of the various chemical and physical means to tune BBB permeability,^6^ focused ultrasound combined with circulating microbubbles has shown potential in achieving localized and reversible BBB opening,^7, 8^ which has entered early-phase clinical investigations.^9–12^

In recent years, light-based approaches have attracted increasing interest in neural stimulation and modulation, because they can achieve spatial resolution down to the single-cell scale.^13–17^ Under short-pulse illumination, when the laser irradiation duration is shorter than the thermal dissipation time, the deposited photon energy causes rapid local expansion and generates transient pressure waves with optical resolution, known as optoacoustic (photoacoustic) effect,^18^ or even inducing cavitation effect under certain circumstances. This has led to ongoing pursuit of exploiting this strong nonlinear light-matter interactions for neural stimulation,^19–22^ and to physically modulate brain vasculature and BBB permeability. In 2011, Choi et al. directed an 800 nm femtosecond laser through a thinned-skull window and focused it on the wall of a selected cortical vein.^23^ This produced transient permeability at single-vessel resolution, probably through cavitation. Subsequently, Yuan et al. used PEGylated gold nanostars to enhance ultrafast laser energy deposition within a multiphoton focal volume,^24^ enabling plasmonic enhancement of brain-tumor microvascular permeabilization. Emelianov and co-workers developed a laser-activated perfluorocarbon nanodroplet strategy for mechanical BBB disruption.^25^ These non-targeted approaches require either high laser intensities or concurrent imaging to localize the vascular distribution of the optoacoustic nanotransducers. More recently, Qin et al. reported mechanobiological modulation of BBB permeability using 532 nm picosecond laser stimulation of gold nanoparticles targeted to endothelial tight junctions.^26, 27^ In their approach, the gold nanoparticles amplify the optoacoustic field and transfer mechanical energy to the molecularly targeted junctions, thereby promoting transient BBB opening and enhanced drug delivery across the blood-brain-tumor barrier in glioblastoma models.^28^

Compared with visible light, near-infrared (NIR) excitation is more attractive for optical modulation of BBB permeability because it provides greater tissue penetration and lower background interference.^29–32^ Moreover, femtosecond excitation can better confine energy deposition in time and space, facilitating impulsive stress generation while restricting thermal diffusion into adjacent tissues and mitigating potential photothermal damage.^33^ To increase local NIR photon absorption and energy deposition, we considered hyperbranched gold nanoconstructs (HBGNCs), whose densely coupled nanoscale branches provide broadband NIR absorption, high absorption efficiency, and strong optoacoustic responses.^34^ Compared with more commonly used gold nanorods and gold nanostars, HBGNCs can maintain strong NIR absorption within a more compact volume.

Here, we developed a strategy for near-infrared reversible optoacoustic modulation of the blood–brain barrier permeability (NIR-ROAMBBB) by combining size-optimized HBGNCs, BV11-mediated endothelial-junction targeting, and spatially confined femtosecond laser excitation. We systematically quantified HBGNCs’ absorption, scattering, and optoacoustic output at 800 nm under matched optical-density conditions, revealed a non-monotonic size dependence and identified 62 nm HBGNCs as the most effective transducers under the conditions examined. We functionalized the selected HBGNCs with BV11 antibodies to recognize junctional adhesion molecule A (JAM-A) at cerebrovascular endothelial junctions, thereby combining molecular localization with spatially confined 800 nm femtosecond laser excitation. We evaluated NIR-ROAMBBB in an endothelial Transwell model and in mice, focusing on its fluence dependence, spatial confinement, reversibility, and capacity to enhance regional accumulation of a model nanoscale cargo. We showed that transient and reversible modulation of BBB permeability with millimeter-scale lateral confinement can be achieved. To the best of our knowledge, this is the first study to combine systematic size optimization of hyperbranched gold optoacoustic transducers with NIR femtosecond excitation for spatially confined and reversible modulation of the intact BBB in vivo. These findings establish the feasibility of NIR plasmon enhanced reversible BBB permeability modulation under femtosecond laser excitation.

## Results and Discussion

### Size tuning of HBGNCs via seeded growth

HBGNCs were synthesized via a secondary seed-mediated strategy, adapted from the nucleation-suppressed growth method reported by Gao et al.,^35^ so that their overall morphology can remain the same despite large-size differences (Figure 1a). Primary Au seeds of 3 nm were first made and then enlarged into approximately 30 nm Au nanospheres, which served as secondary seeds for subsequent HBGNC growth (Figure S1). The existence of PVP stabilized the reduced Au species and limited particle aggregation, while the mild reduction potential of ascorbic acid favored Au deposition onto the pre-existing seeds. This allows size tuning of HBGNCs via the ratio of Au precursor to the concentration of secondary seeds. With the amount of Au precursor and all other growth conditions fixed, increasing the concentration of secondary seeds produced smaller HBGNCs. Conversely, decreasing the seed concentration made more Au available to each seed and yielded larger particles.

**Figure 1.**
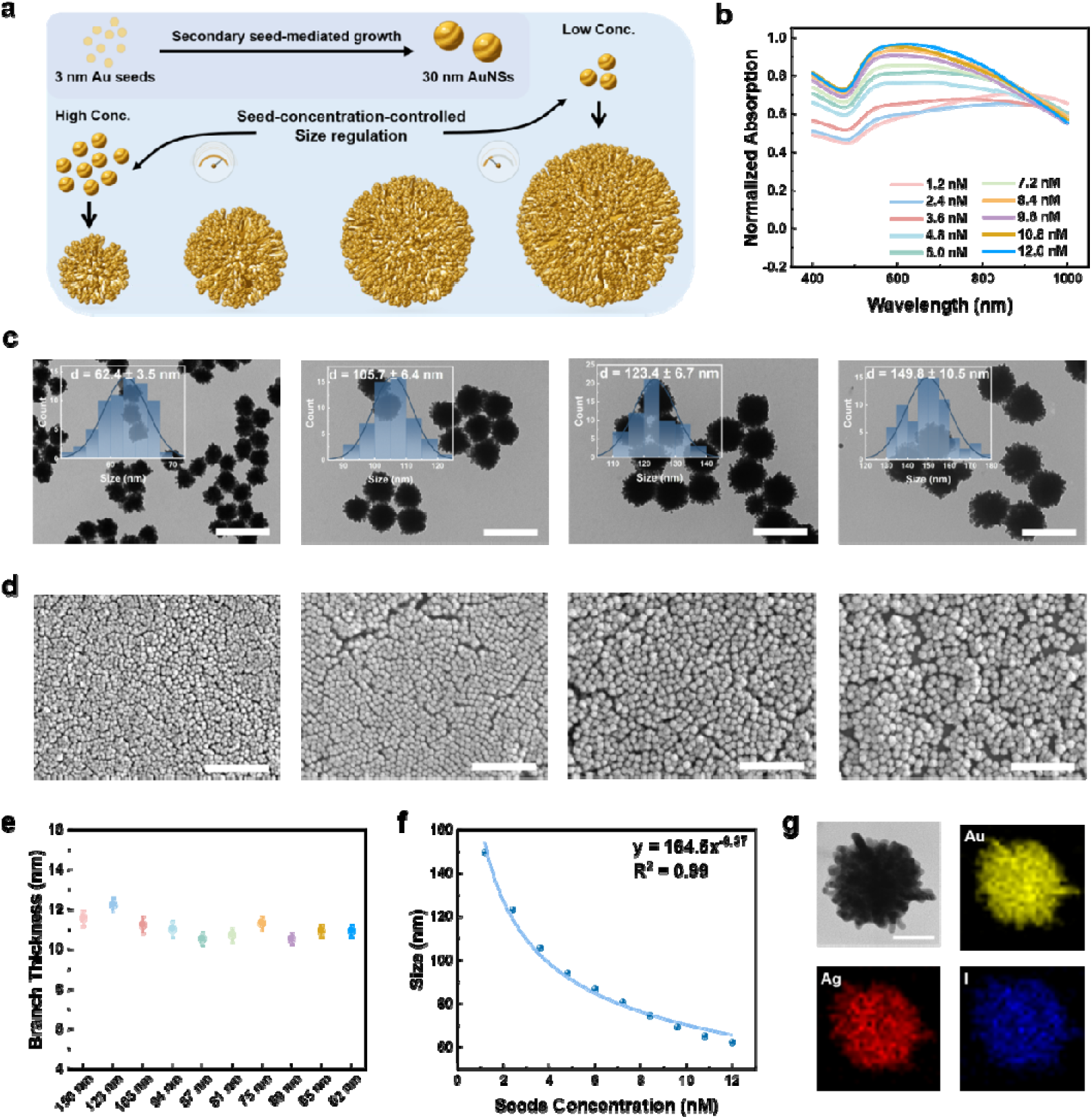
Size-controlled synthesis of HBGNCs. (a) Schematic of HBGNC synthesis using gold nanosphere seeds, with final particle size controlled by the seed concentration. (b) UV-vis-NIR spectra of HBGNCs synthesized at different seed concentrations. (c) Representative high-magnification TEM images and the corresponding particle-size distribution histograms. (d) Representative low-magnification SEM images of HBGNCs of different sizes. (e) Quantification of branch thickness for HBGNCs of different sizes. (f) Fit of HBGNC size as a function of seeds concentration. (g) STEM-EDS elemental maps of HBGNCs. Scale bars, 200 nm (c), 1 μm (d), and 20 nm (g).

To establish the relationship between secondary-seed concentration and final HBGNC size, we systematically varied the concentration of the 30 nm Au seeds while keeping the growth solution composition and other reaction parameters fixed. UV-vis-NIR spectra showed that all HBGNC formulations exhibited broadband extinction across the near-infrared region, with the long-wavelength (>900 nm) extinction slightly strengthened as particle size increased (Figure 1b). Transmission electron microscopy (TEM, Figure 1c and Figure S2) and scanning electron microscopy (SEM, Figure 1d and Figure S3) analyses revealed that all HBGNCs exhibited relatively uniform dimensions, with the mean overall diameter progressively increased from 62.4±3.5 nm to 149.8±10.5 nm with decreasing seed concentrations. Despite the pronounced variation in the overall HBGNC diameters, the mean branch thickness remained at approximately 11 nm across all formulations, with no apparent dependence on HBGNC sizes (Figure 1e). Quantitative analysis showed that the mean HBGNC diameter followed an inverse power-law dependence on the secondary-seed concentration, consistent with seed-number-dependent partitioning of the available Au precursor (Figure 1f). Thus, varying only the secondary seed concentration enabled HBGNC size to be tuned while maintaining a constant growth solution formulation and a common branching mechanism.

In the synthetic mixture, there also existed a relatively low concentration of iodide (∼0.4 mM) and Ag (∼0.2 mM), which served as a surface-regulating species. Ag-modified surface sites and adsorbed iodide generated partially passivated regions on the secondary seeds. This heterogeneous passivation favored Volmer-Weber-type Island deposition over conformal Frank–van der Merwe growth, directing the formation of the hyperbranched architecture.^36^ Scanning transmission electron microscopy-energy dispersive X-ray spectroscopy (STEM-EDS) elemental mapping identified Au as the predominant constituent and showed the presence of Ag and I throughout the branched structures. Inductively coupled plasma mass spectrometry (ICP-MS) analysis of the purified HBGNCs yielded normalized elemental fractions of 89.7% Au, 8.9% Ag and 1.4% I (Figure S4), further confirming the retention of minor but measurable Ag and I components in the final particles. Together with elemental mapping, this composition was consistent with the participation of Ag^+^ and I^-^ containing species in surface-regulated HBGNC growth.

Collectively, these results demonstrate that varying the secondary seed concentration enabled controlled synthesis of HBGNCs over a broad size range, while largely preserving their hyperbranched morphology and characteristic branch dimensions.

### HBGNCs exhibit enhanced and non-monotonic size-dependent optoacoustic responses

Under pulsed laser irradiation, optical energy absorbed by HBGNCs can be rapidly converted into a thermal transient, inducing thermoelastic expansion and generating acoustic waves. We first investigated the optoacoustic (OA) responses of HBGNCs as a function of their sizes. Solutions of HBGNCs with different diameters were loaded into capillary tubes, with each dispersion adjusted to the same optical density at 800 nm (A_800_ = 1). Based on conventional size-dependent optical behavior, a progressive decrease in absorption and hence OA amplitude would be expected as the particle size increases. However, a non-monotonic size dependent OA response for HBGNCs has been observed. The OA amplitude decreased first with increasing HBGNCs size, and then the trend was reversed as the size increased further, with the minimum OA amplitude occurred on HBGNC of 105 nm in diameter (Figure 2a, b). To verify this unusual phenomenon, we also carried out solution photothermal characterization on the same size series under extinction-matched conditions (A_800_ = 1), as photothermal and optoacoustic effects are positively correlated in general. Interestingly, the photothermal measurements reproduced the non-monotonic trend observed in the OA experiments (Figure 2c and Figure S5).

**Figure 2.**
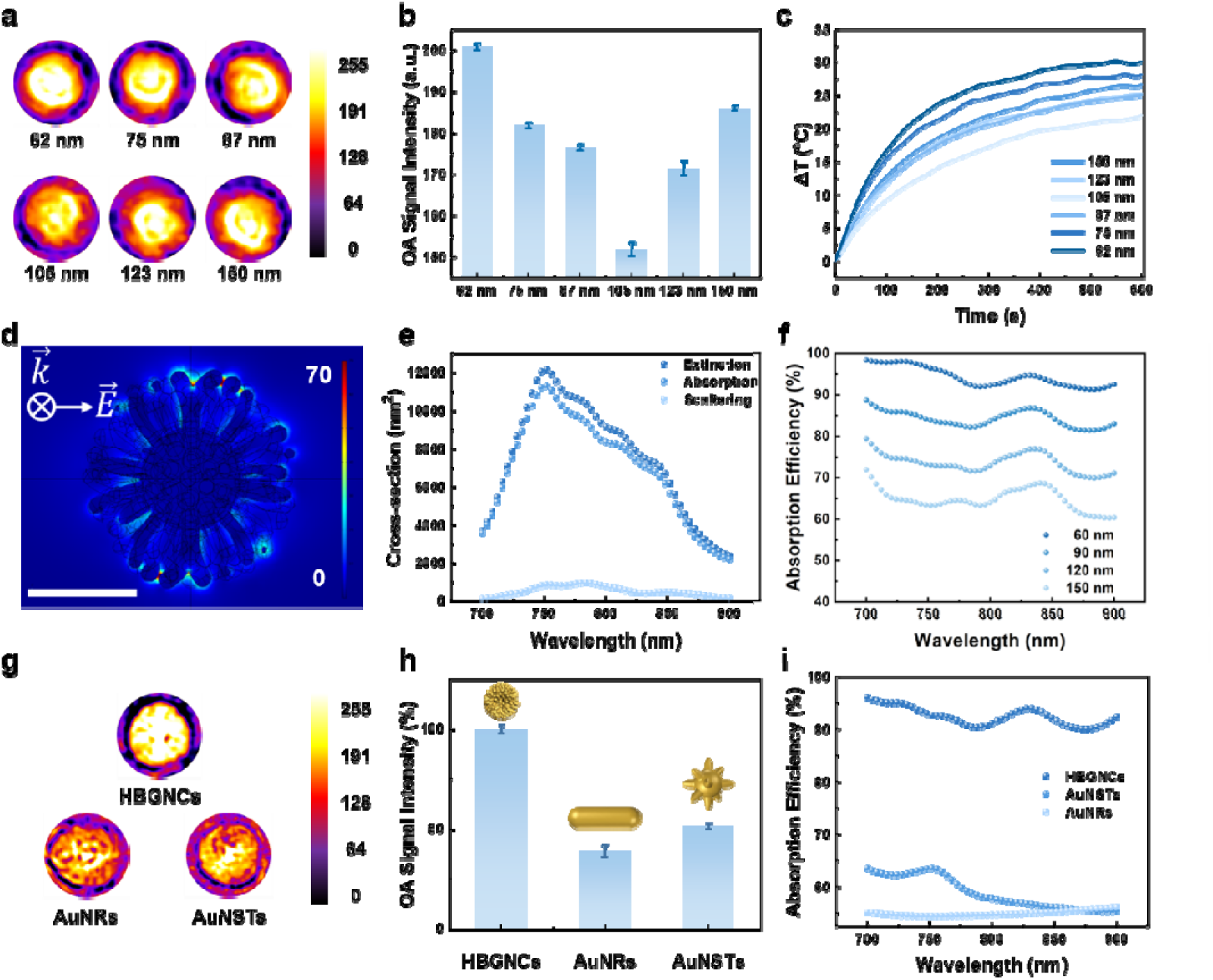
Optoacoustic performance and associated optical properties of HBGNCs. (a) Optoacoustic images of differently sized HBGNCs at matched optical extinction. (b) Optoacoustic signal intensities of differently sized HBGNCs. (c) Photothermal heating (808 nm, 0.75 W/cm^2^) curves of differently sized HBGNCs. (d) Theoretical near-field distribution of HBGNCs at 800 nm. Scale bar, 50 nm. (e) Simulated extinction, absorption, and scattering spectra of HBGNCs over the wavelength range of 700-900 nm. (f) Calculated absorption efficiency of HBGNCs with different sizes over the wavelength range of 700-900 nm. (g) Optoacoustic images of HBGNCs, AuNRs, and AuNSTs. (h) Optoacoustic signal intensities of HBGNCs, AuNRs, and AuNSTs. (i) Calculated absorption efficiencies of HBGNCs, AuNSTs, and AuNRs over the wavelength range of 700-900 nm.

To investigate the plasmonic properties of HBGNCs, we constructed a TEM-informed COMSOL model of the 62 nm HBGNCs and calculated their absorption, scattering, and extinction cross-sections over 700-900 nm. Pronounced local-field enhancement was observed within the inter-branch gaps, consistent with the coupling and hybridization of plasmonic modes supported by adjacent branches (Figure 2d). The absorption efficiency, defined as absorption-to-extinction ratio, reached approximately 95% in the near-infrared region, indicating that most of the extinguished optical energy was dissipated through absorption rather than scattering (Figure 2e). We then calculated the absorption-to-extinction ratios of the different sized HBGNCs using COMSOL Multiphysics. The simulations predicted a monotonic decrease in relative absorption with increasing particle size, accounting for the strong response of the smallest HBGNCs but not for the secondary enhancement observed for the 150 nm particles (Figure 2f). Thus, electromagnetic extinction partitioning alone was insufficient to explain the observed OA trend. Size-dependent heat transfer, interfacial thermal dissipation, and thermoelastic coupling remain possible contributors, which require more thorough investigation in the future.

Next, to benchmark the OA performance of the 62 nm HBGNC against more commonly applied Au nanostructures such as Au nanorods (AuNRs) and gold nanostars (AuNSTs), we loaded the respective solutions into capillary tubes and adjusted their optical density to be the same (A = 1.5) at the excitation wavelength (Figure S6). Under identical irradiation and acquisition conditions, HBGNCs generated the strongest OA response, substantially exceeding those of the reference agents, reaching ∼2 times of that from the AuNRs and AuNSTs (Figure 2g, h). The tendency is in good agreement with the calculated absorption efficiencies, with HBGNCs at approximately 95%, whereas AuNRs and AuNSTs were about 55% and 65%, respectively (Figure 2i and S7). Moreover, the calculated relative absorption of HBGNCs remained nearly invariant across the tested polarization angles, whereas the anisotropic AuNRs exhibited marked orientation-dependent attenuation (Figure S8 and S9). The high surface-to-volume ratio of the small HBGNCs could additionally facilitate rapid heat transfer to the surrounding medium. Collectively, these results identify the densely branched and quasi-isotropic architecture of HBGNCs as an important optical basis for their enhanced PA performance.

### In vitro evaluation of NIR-ROAMBBB

Based on the strongest optoacoustic output, efficient thermal response and smaller hydrodynamic size, the 62 nm HBGNC was selected for subsequent biological investigations. To specifically target the brain endothelial tight junctions, we functionalized HBGNCs surfaces with BV11, an antibody against junctional adhesion molecule A (JAM-A). BV11 was first coupled to OPSS-PEG-SVA and then immobilized on HBGNCs, after which residual surface sites were passivated with mPEG (Figure 3a). After BV11 functionalization, the near-infrared extinction band redshifted slightly (Figure 3b), the hydrodynamic diameter increased from 110 to 135 nm, and the ζ-potential became more negative (from +17.5 mV to -26.5 mV, Figure S10). Together, these spectral and colloidal changes indicated the successful BV11 conjugation on HBGNCs.

**Figure 3.**
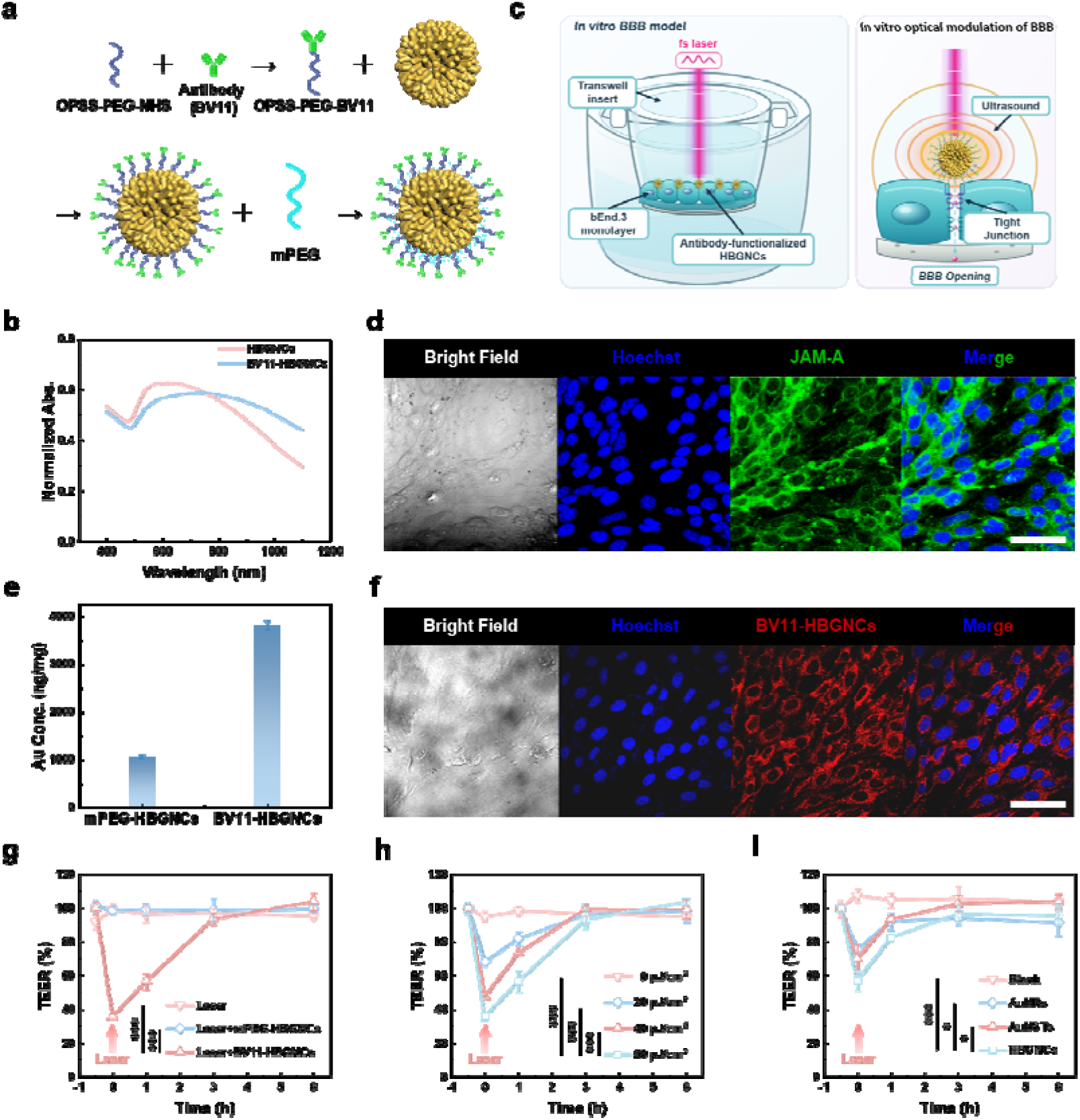
In vitro evaluation of femtosecond-laser-mediated reversible modulation of blood-brain barrier (BBB) permeability. (a) Preparation of BV11-functionalized HBGNCs (BV11-HBGNCs). (b) UV-vis-NIR spectra of BV11-HBGNCs and HBGNCs. (c) Schematic of the in vitro BBB model and BBB opening mediated by femtosecond-laser-induced optoacoustic effects. (d) Bright field, nuclei and tight-junction protein JAM-A fluorescence images of the cell monolayer. (e) Inductively coupled plasma mass spectrometry (ICP-MS) analysis of Au contents following incubation of non-targeting and targeting HBGNCs with the cell monolayer. (f) Bright field, nuclei and BV11-HBGNCs fluorescence images of the cell monolayer. (g-i), Trans-endothelial electrical resistance (TEER) measurements of changes in BBB permeability following femtosecond laser irradiation. (g) TEER changes in the control and experimental groups. (h) Temporal evolution of TEER as a function of the laser fluence. (i) Comparison of BBB-opening efficacy mediated by HBGNCs, AuNRs, and AuNSTs. Scale bar, 50 μm. Data are expressed as mean ± S.D. (n = 3): *, P < 0.05, **, P < 0.01; ***, P < 0.001.

For in vitro studies, we first established a blood-brain barrier model by culturing murine brain endothelial bEnd.3 cells to confluence on Transwell inserts (Figure 3c). The resulting monolayers reached trans-endothelial electrical resistance (TEER) values ∼30 Ω cm² after 4 days of culturing (Figure S11). In addition, immunofluorescence staining showed continuous junctional expression of JAM-A (Figure 3d), validating the formation of a confluent endothelial barrier with organized tight junctions. After being incubated with BV11-HBGNC, ICP-MS analysis revealed 4-fold more cell-associated Au than that from a non-targeting mPEG-HBGNC counterpart (Figure 3e), demonstrating effective molecular targeting of BV11-functionalized HBGNC to the endothelial layer.

To better visualize the specific targeting of BV11-HBGNCs, we prepared a biotinylated tracer by replacing the mPEG backfilling ligand with biotin-PEG-SH, which can then be detected using fluorescent streptavidin. Following incubation with bEnd.3 monolayers, the fluorescence image showed a reticular pattern along the boundaries between neighboring cells (Figure 3f), consistent with preferential nanoparticle accumulation at the endothelial cell–cell junctions. Thus, fluorescence microscopy imaging established the spatial enrichment of BV11-HBGNCs at intercellular junctions, whereas elemental analysis independently quantified their increased association with the endothelial monolayer. Collectively, these results supported BV11-mediated targeting of HBGNCs towards JAM-A-enriched endothelial junctions.

We next evaluated NIR-ROAMBBB in the endothelial Transwell model by monitoring normalized TEER (Figure 3g) after 30 s femtosecond laser irradiation (800 nm, 95 fs, 12.5 kHz). Laser irradiation alone or irradiation in the presence of non-targeting mPEG-HBGNCs left TEER close to baseline, remained at approximate 108% and 97%, respectively. By contrast, the TEER fell to approximately 35% of the original value 5 min after irradiation in the BV11-HBGNC group. TEER in the BV11-HBGNC group subsequently recovered to approximately 56% at 1 h and 93% at 3 h post laser irradiation, and eventually returned to baseline by 6 h. These results showed that BV11 functionalization was necessary for the pronounced and reversible barrier opening under the tested conditions. In addition, we evaluated the phototoxicity of the femtosecond laser and the effect of HBGNCs on cell viability, and the results demonstrated that after laser irradiation alone and incubation of BV11-HBGNCs for laser irradiation, the cells all maintained viability over 85% (Figure S12), suggesting excellent biocompatibility of HBGNCs.

The magnitude of the TEER response also depends on the applied laser fluence (Figure 3h). TEER declined to approximately 68%, 48%, and 35% of the baseline value after being irradiated at 20, 40, and 60 μJ/cm², respectively. Despite the different degrees of initial TEER reduction, they all recovered and approached the original value within 3–6 h post femtosecond laser irradiation. The fluence dependence and subsequent recovery demonstrated the tunable, transient, and reversible modulation of the brain endothelial barrier.

To demonstrate the advantage of applying HBGNCs over commonly used AuNRs and AuNSTs, we have incubated the respective solutions with the same optical density over the bEnd.3 monolayers, followed by the same femtosecond laser irradiation treatment (Figure 3i). As expected, HBGNCs produced the largest immediate decrease in TEER, as opposed to AuNRs and AuNSTs. Together, these results identified BV11-mediated junctional targeting, laser fluence, and nanoparticle formulation as three determinants of the measured barrier response.

### In vivo evaluation of NIR-ROAMBBB

Encouraged by the promising in vitro performance, we then went on to assess NIR-ROAMBBB in vivo by examining cerebral HBGNC accumulation, Evans blue extravasation, permeability recovery, and the spatial geometry of the response (Figure 4a). We acquired transcranial optoacoustic images under identical conditions at 1 h post intravenous injection of BV11-HBGNCs or mPEG-HBGNCs, in which BV11-HBGNCs produced a higher OA signal within the brain region of interest than the non-targeted mPEG-HBGNCs (Figure 4b). Independently, ICP-MS analysis revealed that the brain-associated gold (Au) content in the BV11-HBGNCs group was approximately four times higher than that in the mPEG-HBGNCs group (Figure S13). These concordant measurements demonstrated that BV11 functionalization increased the cerebral enrichment of HBGNCs.

**Figure 4.**
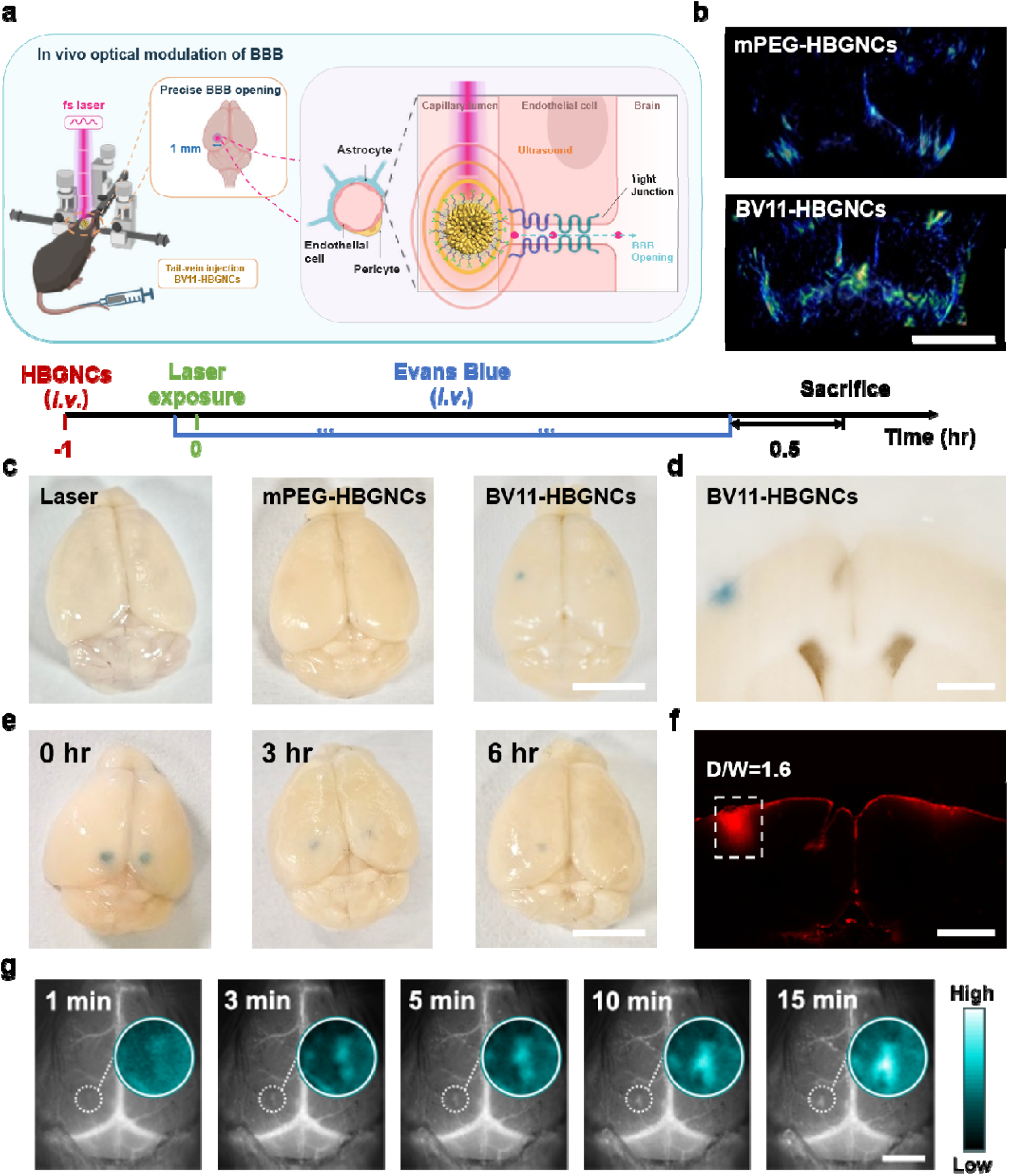
In vivo reversible modulation of blood-brain barrier permeability using tight-junction-targeting HBGNCs and femtosecond laser irradiation. (a) Schematic of transcranial femtosecond laser irradiation combined with BV11-HBGNCs to modulate blood-brain barrier (BBB) permeability in mice. (b) Optoacoustic images of mouse brain following i.v. injection of mPEG-HBGNCs and BV11-HBGNCs. (c) Representative pictures of mouse brain from 3 groups after the outlined experimental procedures. Laser: i.v. injected PBS; mPEG-HBGNCs and BV11-HBGNCs refer to the i.v. injected substances, laser fluence 0.25 mJ/cm^2^ and a spot diameter of 1 mm. (d) Photograph of brain section showing Evans blue extravasation from the BV11-HBGNCs treatment group. (e) Assessment of the reversibility of BBB permeability modulation. The denoted time refers to the point when Evans blue was systemic administered after laser irradiation. The left and right sides were irradiated at fluences of 1 and 0.25 mJ/cm², respectively. (f) Wide-field inverted fluorescence microscopy image of brain section following Evans blue extravasation, together with quantification of the leakage depth (D) and width (W). (g) In vivo fluorescence imaging of the accumulation of a model delivery cargo, mPEG-AgAuSe quantum dots, over the irradiated brain region. Scale bars, 5 mm (b, c and e) and 2.5 mm (d, f and g). Data are expressed as mean ± S.D. (n = 3 mice).

We next examined whether the increased cerebral enrichment could support focal modulation of BBB permeability. One hour after tail vein administration of BV11-HBGNCs (18.9 µg/g), a selected cortical region was transcranially irradiated with an 800 nm femtosecond laser (95 fs, 12.5 kHz, 1 mm beam diameter) for 120 s to induce transient local BBB opening. The permeability of the blood-brain barrier was evaluated by Evans blue-albumin extravasation, where Evans blue was injected through the tail vein and allowed to circulate for half an hour. Focal Evans blue extravasation was detected within the irradiated region, indicating transient passage of the tracer from the vasculature into the surrounding brain tissue (Figure 4c, d). Neither femtosecond irradiation alone nor mPEG-HBGNCs combined with identical irradiation produced detectable focal extravasation. Thus, laser exposure alone and non-targeted HBGNCs under the same irradiation conditions were insufficient to reproduce the permeability increase observed with BV11-HBGNCs. We also found that the duration of the permeability window depended on laser fluence. At the lower fluence condition (0.25 mJ/cm^2^), Evans blue-detectable permeability returned to the baseline range within 3 h after irradiation. While at 1 mJ/cm^2^, Evans blue extravasation persisted longer, and approached baseline levels by approximately 6 h (Figure 4e). These results defined a fluence-dependent permeability window that remained transient under both irradiation conditions. Brain sections collected at 30 min after treatment were examined by H&E staining. The irradiated cortical region retained an overall architecture comparable to that of the contralateral region, with no overt cavitation, extensive hemorrhage, or widespread histomorphological disruption under the tested conditions (Figure S14).

We then quantified the spatial geometry of the permeability response using wide-field fluorescence microscopy of brain sections. Evans blue-positive regions were segmented using a consistent fluorescence threshold, from which the axial leakage depth and lateral width were determined. The leakage reached a mean depth of 1.6 mm but remained confined to a mean lateral width of 1.0 mm. This yielded a depth-to-width ratio of approximately 1.6, with the axial extent exceeding the lateral spread by approximately 60% (Figure 4f). Results reported in previous studies yielded depth-to-width ratios of only 0.5–0.6.^26, 27^ Although direct cross-study comparison is limited by differences in experimental and imaging conditions, this contrast is consistent with preferential axial extension and limited lateral spread under our experimental conditions. We speculate this can be due to the combined effect of better NIR penetration and more confined excitation from femtosecond laser.

Finally, we investigated whether the transient increased permeability window could promote the regional delivery of nanoscale cargo. We utilized NIR-II emitting mPEG functionalized AgAuSe quantum dots (mPEG-AgAuSe QDs) as a model cargo for real-time visual inspection (Figure S15), which were administered intravenously after five minutes of laser irradiation. Using 808 nm laser excitation, time resolved fluorescence imaging showed a rapid increase in intensity within the laser treated brain region (Figure 4g), and the regional fluorescence reached its maximum approximately 15 min after administration and subsequently plateaued, demonstrating rapid enrichment of the nanoscale cargo at the site of BBB modulation. Collectively, these findings show that cerebral enrichment of BV11-HBGNCs enabled spatially confined, reversible and fluence tunable modulation of BBB permeability in vivo.

Under current experimental settings, we do not have a direct way to measure the local temperature rise in situ. Therefore, we examined the optical stability and thermal response of 62 nm HBGNCs solutions under similar laser parameters used in the animal experiments. 1 mL HBGNC dispersion with an optical density of 3.5 at 800 nm (A_800_ = 3.5) was exposed to femtosecond-laser irradiation for 2 min. The extinction spectrum remained essentially unchanged after irradiation, indicating no detectable alteration in the ensemble optical response of the HBGNCs (Figure S16). Infrared thermography and time-resolved temperature measurements showed a bulk solution temperature increase of approximately 5 °C over the same period (Figure S17 and S18). Considering the high optical loading used in this solution-phase stress test, these results suggest that substantial HBGNC-mediated bulk heating is unlikely under the lower particle burden expected in the brain for in vivo studies.

## Conclusion

In summary, we developed NIR-ROAMBBB by integrating size-optimized HBGNCs, BV11-mediated endothelial-junction targeting, and 800 nm femtosecond laser excitation. Systematic size screening identified 62 nm HBGNCs as the most efficient optoacoustic transducer under 800 nm excitation, which yielded approximately two-fold stronger optoacoustic signal of conventional gold nanorods and nanostars. Surface functionalization with BV11 efficiently anchored HBGNCs to JAM-A-enriched brain endothelial junctions, enabling transient BBB permeability modulation under synergistic femtosecond pulsed laser excitation, which is not feasible under either laser irradiation alone or with non-targeted HBGNCs under identical irradiation conditions. The induced permeability enhancement was tunable via laser fluence and spontaneously reversed back to baseline levels within hours in both in vitro cell models and in vivo mouse models. This defined permeability window enabled the localized accumulation of a model nanoscale cargo in brain tissues. The demonstrated NIR-ROAMBBB strategy offers a novel strategy for precise, localized, and reversible blood–brain barrier opening for brain delivery applications.

