## Supplemental Information for "Near-infrared optoacoustic modulation of the blood-brain barrier permeability using size-tuned hyperbranched gold nanoconstructs"

### Materials

Cetyltrimethylammonium bromide (CTAB, 99%), Anti-JAM-A Antibody (clone BV11), fluorescein isothiocyanate-dextran (FITC-dextran, 40 kDa), Evans blue, and polyvinylpyrrolidone (PVP, molecular weight 55k) were purchased from Sigma-Aldrich. Silver nitrate ($AgNO_3$, 99%), chloroauric acid ($HAuCl_4 \cdot 4H_2O$), hydrochloric acid (HCl, 36-38%), ascorbic acid (AA, 99.7%), potassium iodide (KI), and sodium borohydride ($NaBH_4$) were purchased from Sinopharm Chemical Reagent Co., Ltd. Sodium oleate (NaOL, 98%) was purchased from TCI. Methoxy polyethylene glycol thiol (mPEG2000-SH) was purchased from Xi'an Ruixi Biological Technology. Biotin-PEG1000-SH was purchased from Pengshuo Biotechnology. Orthopyridyl disulfide-PEG-succinimidyl valerate (OPSS-PEG3500-SVA) was purchased from Laysan Bio, Inc. DMEM and fetal bovine serum (FBS) were purchased from Gibco. Penicillin-Streptomycin solution, Hoechst 33342, boric acid buffer solution (0.1 M, pH 8.5), 4% paraformaldehyde (PFA), (3-(4,5-dimethylthiazol-2-yl)-2,5-diphenyl tetrazolium bromide (MTT), and dimethyl sulfoxide (DMSO) were purchased from Beyotime.

### Synthesis of HBGNCs

3 nm gold seeds were prepared by direct $NaBH_4$ reduction, with sodium citrate serving as the capping agent.[1] 1 mL of 5 mM $HAuCl_4$ and 1 mL of 5 mM sodium citrate were added to 18 mL of deionized water. While the mixture was stirred vigorously, 0.6 mL of freshly prepared 0.1 M $NaBH_4$ was rapidly injected, after which the solution turned reddish yellow. The resulting dispersion was stirred at room temperature for at least 4 h. It was then collected and stored at 4 °C in the dark until use as a seed stock for subsequent gold nanospheres (AuNSs) growth.

30 nm AuNSs were synthesized from the 3 nm gold seeds by seed-mediated growth,[2] with the growth reactions performed in 20 mL scintillation vials. Each vial received 0.5 mL of 5 wt% PVP, 0.25 mL of 0.1 M ascorbic acid, and 0.2 mL of 0.2 M KI. Next, 1.6 mL of 10 mM $HAuCl_4$ and 0.46 mL of deionized water were added. The mixture was then stirred at 900 rpm for 1 min. Under vigorous stirring, 170 μL of the 3 nm seed solution was rapidly injected. The reaction mixture was incubated without

agitation at 30 °C for 10 min. The product was purified three times by centrifugation at 10,000 rpm for 15 min per cycle. The collected AuNSs were redispersed in water, adjusted to 12.0 nM and stored at 4 °C in the dark until use.

HBGNCs were then synthesized from 30 nm AuNSs by partial surface passivation.[3] To prepare the growth solution, 100 μL of 10 wt% PVP and 250 μL of a 5 mM $AgNO_3$ solution were added to 4 mL of deionized water. The mixture was magnetically stirred at 900 rpm for 5 min to ensure homogeneity. For size control, seed solutions of different concentrations were prepared from the 12.0 nM AuNS stock. Under vigorous stirring, 200 μL of the selected 30 nm AuNS seed solution was added, followed by 100 μL of 20 mM KI and 288 μL of 21.718 mM $HAuCl_4 \cdot 4H_2O$. Finally, 200 μL of 0.5 M ascorbic acid was rapidly injected, causing the solution to change rapidly from light pink to black. The reaction was maintained at 30 °C for 30 min. HBGNCs were washed three times with deionized water by centrifugation at 9,000 rpm for 15 min per cycle. The collected HBGNCs were redispersed in deionized water and stored at 4 °C until further use.

### Synthesis of Gold Nanostars (AuNSTs)

Gold nanosphere seeds with diameter of 12 nm were prepared by citrate reduction. 10 mL of 1 mM $HAuCl_4$ was brought to boiling in an oil bath. Next, 1.5 mL of 1% sodium citrate solution was rapidly injected, and the reaction mixture was boiled for 15 min. After cooling to room temperature, the seed dispersion was filtered through a 0.22 μm nitrocellulose membrane. The absorbance at 520 nm ($A_{520}$) was adjusted to 2.61, and the seed dispersion was stored at 4 °C until use.

AuNSTs were synthesized by seed-mediated growth at 25 °C. In a 20 mL scintillation vial, 10 mL of 0.25 mM $HAuCl_4$ was mixed thoroughly with 10 μL of 1 M HCl. Next, 75 μL of citrate-stabilized seed dispersion was added. While stirring at 700 rpm, 100 μL of 4 mM $AgNO_3$ and 50 μL of 100 mM ascorbic acid were rapidly injected simultaneously. After 30 s of stirring, the solution changed from pale red to blue-green. At this point, 100 μL of 10 wt% PVP was added, and the reaction was stirred for another 2 min. The AuNSTs were washed three times in 15 mL centrifuge tubes by centrifugation at 3,000 g for 15 min to remove excess PVP. The collected product was resuspended in deionized water for subsequent use.[4]

### Synthesis of Gold Nanorods (AuNRs)

The gold seeds were freshly prepared before each AuNR synthesis by $NaBH_4$ reduction. In a 20 mL scintillation vial, 5 mL of 0.5 mM $HAuCl_4$ was mixed with 5 mL of 0.2 M CTAB. Freshly prepared $NaBH_4$ solution (0.6 mL, 10 mM) was diluted to 1 mL with deionized water and injected into the precursor mixture under vigorous stirring. The solution changed from golden yellow to brownish yellow. Stirring was continued for 2 min, and the seed dispersion was then aged at 25 °C for at least 30 min.

In a 100 mL Erlenmeyer flask, 0.9 g of CTAB and 0.1234 g of NaOL were dissolved in 25 mL of water preheated to 60 °C. After the solution was cooled to 30 °C, 2.4 mL of 4 mM $AgNO_3$ was added with stirring. The mixture was held at 30 °C for 15 min. Next, 25 mL of 1 mM $HAuCl_4$ was added, and the solution was stirred at 700 rpm for 90 min until it became colorless. 1.5 mL of concentrated HCl was then added to adjust the solution pH to 1.5. The solution was stirred at 400 rpm for 15 min. Then, 125 μL of 64 mM ascorbic acid was added, followed by vigorous stirring for 30 s. Finally, 20 μL of freshly prepared seed dispersion was injected into the growth solution. After 30 s of stirring, the mixture was incubated without agitation at 30 °C overnight. The AuNRs were washed three times with deionized water by centrifugation at 7,000 rpm for 30 min per cycle. The purified AuNRs were resuspended in deionized water for subsequent use.[5]

### Surface functionalization of HBGNCs

Anti-JAM-A Antibody (BV11) was diluted to 0.05 mg/mL in 2 mM borate buffer (pH 8.5). OPSS-PEG-SVA was dissolved in the same buffer and added to the diluted antibody at an OPSS-PEG-SVA-to-antibody molar ratio of 125:1. The mixture was gently agitated for 4 h on ice in the dark. Unreacted OPSS-PEG-SVA was removed by dialysis against borate buffer for 12 h at 4 °C in the dark using 20 kDa MWCO membrane. The resulting OPSS-PEG-SVA-functionalized antibody was mixed with HBGNCs at an antibody-to-HBGNC molar ratio of 200:1. Conjugation proceeded for

1 h under gentle agitation on ice in the dark. To passivate the remaining HBGNC surface, mPEG-thiol (mPEG-SH; molecular weight 2 kDa) was added at a nominal density of 16 PEG/nm$^2$. The mixture was agitated for 1 h on ice in the dark. The resulting BV11-HBGNCs were washed three times with borate buffer by centrifugation at 9,000 rpm for 15 min at 4 °C. The final product was resuspended in borate buffer and stored at 4 °C in the dark for up to two weeks.[6]

### Characterization

The morphology and structure information were examined by transmission electron microscopy (TEM, Hitachi HT7700) and scanning electron microscopy (SEM, Quanta 250FEG) at an operating voltage of 120 kV and 5 kV, respectively. Scanning Transmission Electron Microscopy-Energy Dispersive X-ray Spectroscopy (STEM-EDS) elemental mapping analysis was performed using a field-emission transmission electron microscope (Tecnai G2 F20 S-TWIN). Extinction spectra were recorded using a UV-vis-NIR spectrophotometer (Shimadzu, UV-3600 Plus). The photoacoustic signal intensity of HBGNCs, AuNRs, and AuNSTs were acquired using PA imaging system (TomoWave, LOIS 3D) equipped with a nanosecond-pulsed laser. All samples were dispersed in water and matched for optical absorbance and suspension volume. The photothermal response of HBGNCs was evaluated by monitoring temperature changes with an infrared thermal camera (Fluke, Ti 300) under fs or continuous wave laser excitation. Zeta potential and hydrodynamic diameter of the nanomaterial dispersions were measured using Dynamic Light Scattering (DLS, Malvern Instruments).

### Numerical simulation

Optical simulations were performed in COMSOL Multiphysics using the Electromagnetic Waves, Frequency Domain interface. Extinction, absorption and scattering cross-sections were calculated for HBGNCs, AuNRs, and AuNSTs. The surrounding medium was set to water at 293.15 K, matching the aqueous conditions used for photoacoustic and extinction-spectroscopy measurements. The wavelength-dependent optical constants of gold were taken from the data reported by Johnson and Christy,[7] which spans from 0.188 to 1.937 μm. A scattered-field formulation was used, with a background plane wave as the excitation source. Spectra were calculated from 700 to 900 nm to obtain the extinction, scattering and absorption cross-sections, together with the absorption efficiency. The structural parameters of each nanoparticle model were derived from the corresponding transmission electron microscopy images. Each simulation represented an isolated nanoparticle.

### In vitro BBB model construction and evaluation

An in vitro blood-brain barrier (BBB) model was established using mouse brain microvascular endothelial (bEnd.3) cells cultured on Transwell inserts. Cells were maintained in DMEM supplemented with 10% FBS and 1% penicillin-streptomycin. After two passages, cells were seeded onto matrix-coated Transwell inserts and cultured for 3-4 days at 37 °C in 5% $CO_2$ to form confluent monolayers. Monolayer integrity was assessed by measuring transendothelial electrical resistance (TEER) with a voltammeter. After subtraction of the blank-insert baseline, monolayers with TEER values exceeding 30 Ω $cm^2$ were used for subsequent experiments. Expression of the tight-junction (TJ) protein JAM-A was confirmed by immunocytochemistry. Monolayers were fixed in ice-cold 4% PFA for 5 min and washed three times with PBS. Cells were blocked for 1 h at room temperature and incubated overnight at 4 °C with 5 μg/mL anti-JAM-A Antibody (BV11). After three PBS washes, cells were incubated with secondary antibody for 1 h at room temperature. Nuclei were stained with Hoechst dye for 10 min in the dark. Fluorescence images were acquired using a confocal microscope (Olympus, FV3000).

### Evaluation of the targeting capability of functionalized HBGNCs

The specific targeting capability of functionalized HBGNCs was evaluated by in vitro fluorescence imaging and elemental quantification method. To enable fluorescence detection, mPEG-thiol was replaced by biotin-PEG-SH during the final backfilling step of BV11-HBGNC conjugation, applied with the same nominal surface density and reaction conditions as those for mPEG-thiol. Using the in vitro BBB model, BV11-HBGNCs-biotin was incubated with bEnd.3 cell monolayers for 30 min at 37 °C under 5% $CO_2$. Then, the monolayers were washed three times with PBS to remove physically attached nanoparticles and fixed with 4% PFA for 10 min at 4 °C. Visualization of BV11-HBGNCs-biotin across the bEnd.3 monolayer was realized by staining with Cy3-labeled streptavidin (diluted 1:200), followed by fluorescence imaging using a confocal microscope (Olympus, FV3000). Immediately after imaging, 1 mL of freshly prepared aqua regia was added directly to each culture dish to digest the sample. The digested solution was diluted and sent to inductively coupled plasma mass spectrometry (ICP-MS) for the elemental analysis. The measured cell-associated gold content was used to assess the quantitative targeting efficiency of BV11-HBGNCs.

### In vitro optical modulation of BBB

After model validation, baseline TEER was recorded. BV11-HBGNCs (0.04 nM) were incubated with the endothelial monolayers for 30 min at 37 °C in 5% $CO_2$. The monolayers were washed three times with PBS. The in vitro BBB models were then irradiated with a femtosecond (fs) laser (800 nm, 95 fs, 12.5 kHz, irradiation for 30 s). TEER was measured at 5 min and 1, 3, and 6 h after irradiation to characterize the time course of barrier opening.

### In vivo biodistribution of HBGNCs

C57BL/6 mice were anesthetized by intraperitoneal injection of 150 μL pentobarbital solution (1 mg/mL). Then, BV11-HBGNCs were administered through the tail vein at 18.9 μg/g body weight. After 1 h of circulation, the brain was imaged using a small-animal photoacoustic imaging system. Relative cerebral accumulation of BV11-HBGNCs was assessed from differences in photoacoustic signal intensity within predefined brain regions.

For quantitative biodistribution analysis, a separate cohort received BV11-HBGNCs by tail-vein injection and was allowed to circulate for 1 h. Deeply anesthetized mice were transcardially perfused with 30 mL of ice-cold PBS to remove residual intravascular blood. The brains were collected, weighed, and microwave-digested, with their gold content quantified by ICP-MS.

### In vivo optical modulation of BBB

C57BL/6 mice were used to assess transcranial femtosecond-laser modulation of

BBB permeability in vivo. Mice were anaesthetized by intraperitoneal injection of 150 μL pentobarbital solution (1 mg/mL). JAM-A-targeting BV11-HBGNCs were administered through the tail vein at 18.9 μg/g body weight. After 1 h of circulation, the scalp was incised to expose the skull. Vascular HBGNCs were irradiated transcranially with a fs laser (800 nm, 95 fs, 12.5 kHz, irradiation for 120 s) through the intact skull with a femtosecond laser. The repetition rate was fixed at 12.5 kHz, and the incident fluence was set to 0.25 or 1.0 mJ/cm$^2$ by adjusting the pulse energy to produce different degrees of BBB opening. At the specified time points after irradiation, 100 μL of 2% Evans blue solution was administered through the tail vein. After 30 min, mice were transcardially perfused with 40 mL ice-cold phosphate-buffered saline to remove intravascular dye. Tissues were subsequently fixed by perfusion with 20 mL 4% paraformaldehyde. Fixed brains were cryosectioned at a thickness of 20 μm at -20 °C. Sections were mounted on glass slides and imaged using a wide-field inverted fluorescence microscope (Leica, DMi8).

Following BBB opening, PEGylated AgAuSe quantum dots (mPEG-AgAuSe QDs; 0.2 mg/mL, 100 μL) prepared following previous protocols were administered intravenously via the tail vein.[8] In vivo fluorescence imaging was subsequently performed under 808 nm laser excitation, and cerebral fluorescence was monitored over time to track the accumulation of mPEG-AgAuSe QDs in the brain parenchyma.

### Supplementary Figures

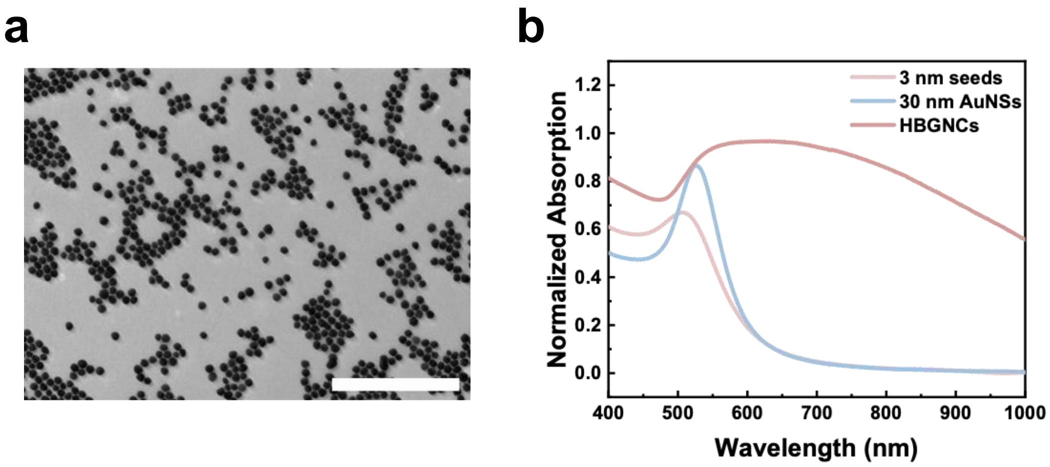


**Figure S1. (**a) TEM image of 30 nm gold nanospheres (AuNSs), scale bar: 500 nm. (b) UV-vis-NIR spectra of 3 nm seeds, 30 nm AuNSs, and HBGNCs.

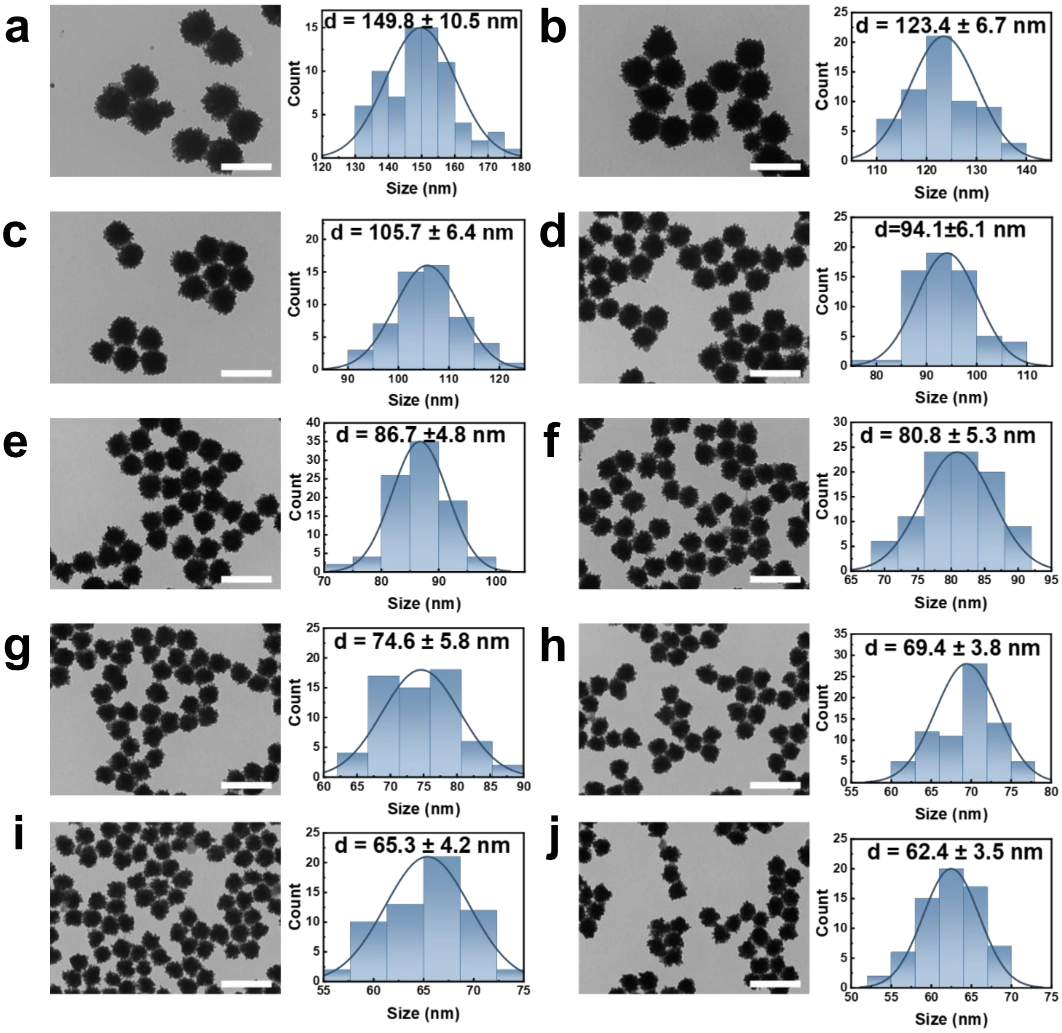


**Figure S2.** TEM images and size distribution histograms of HBGNCs synthesized using different concentration of AuNSs seeds, with (a) 1.2 nM, (b) 2.4 nM, (c) 3.6 nM, (d) 4.8 nM, (e) 6.0 nM, (f) 7.2 nM, (g) 8.4 nM, (h) 9.6 nM, (i) 10.8 nM, and (j) 12.0 nM, respectively. Scale bars, 200 nm.

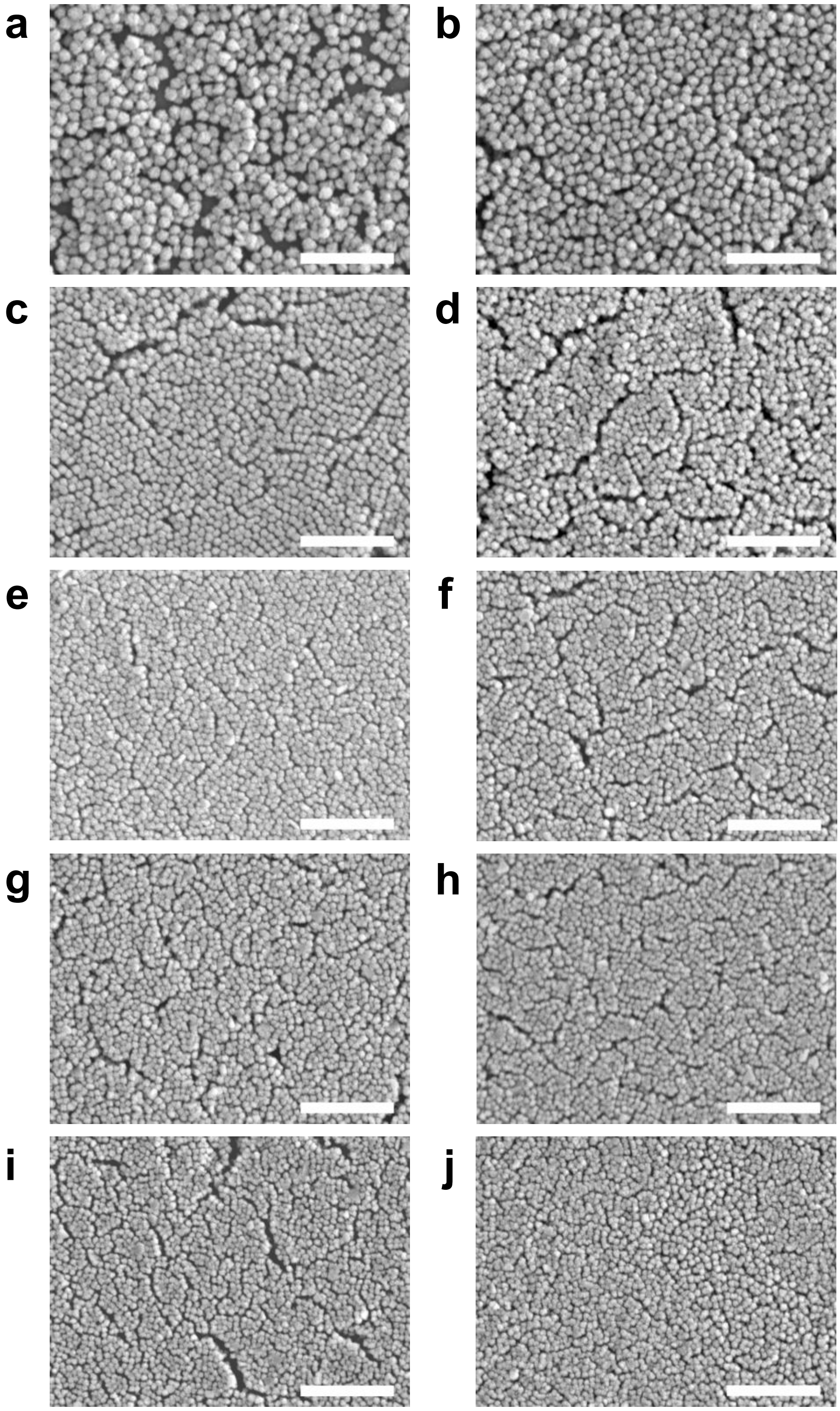


**Figure S3.** SEM images of HBGNCs synthesized using different concentration of AuNSs seeds, with (a) 1.2 nM, (b) 2.4 nM, (c) 3.6 nM, (d) 4.8 nM, (e) 6.0 nM, (f) 7.2 nM, (g) 8.4 nM, (h) 9.6 nM, (i) 10.8 nM, and (j) 12.0 nM, respectively. Scale bars, 1 μm.

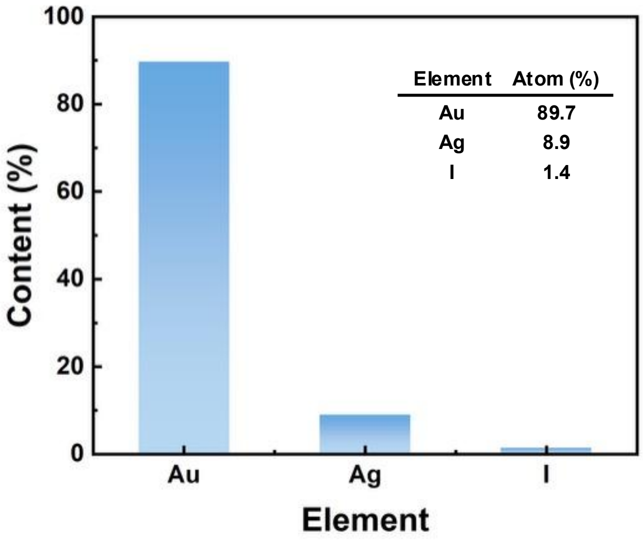


**Figure S4.** Inductively coupled plasma mass spectrometry (ICP-MS) analysis of HBGNCs.

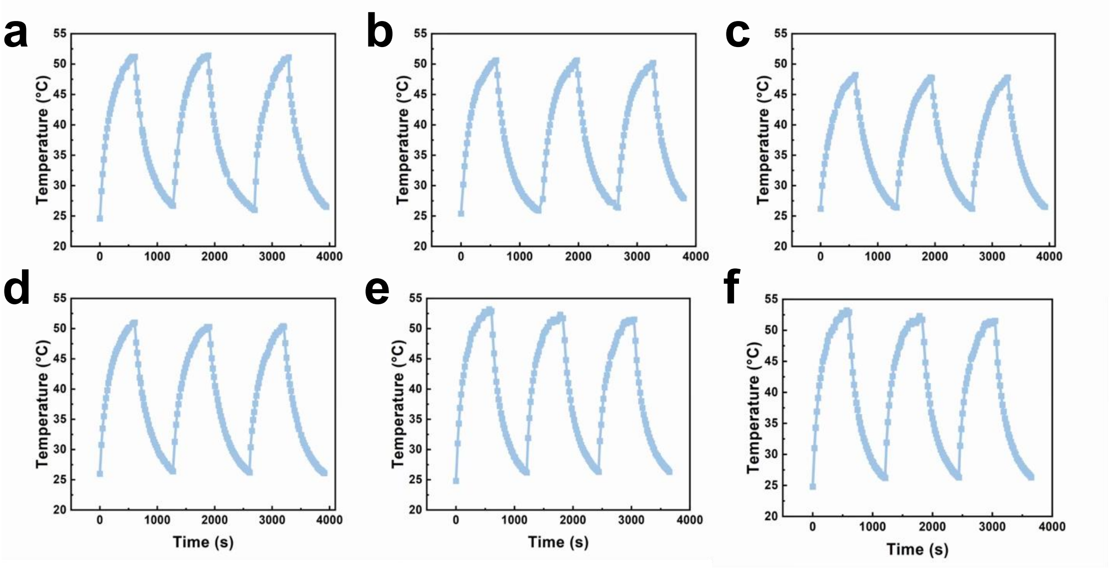


**Figure S5.** Repeated photothermal heating and cooling curves for different sized HBGNCs. (a) 150 nm, (b) 123 nm, (c) 105 nm, (d) 87 nm, (e) 75 nm, (f) 62 nm. Solution was irradiated by 808 nm continuous wave laser at a power density of 0.75 W/cm$^2$.

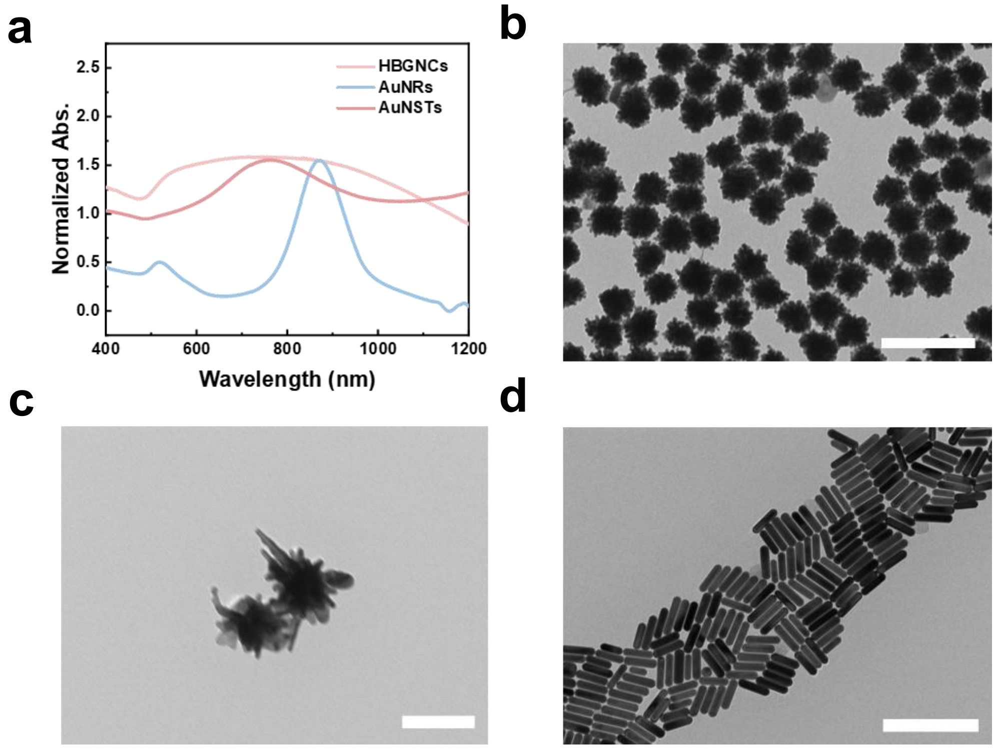


**Figure S6.** (a) UV-vis-NIR spectra of HBGNCs, AuNSTs and AuNRs. (b) TEM images of HBGNCs. (c) TEM images of AuNSTs. (d) TEM images of AuNRs. Scale bars, 200 nm (b and d), 100 nm (c).

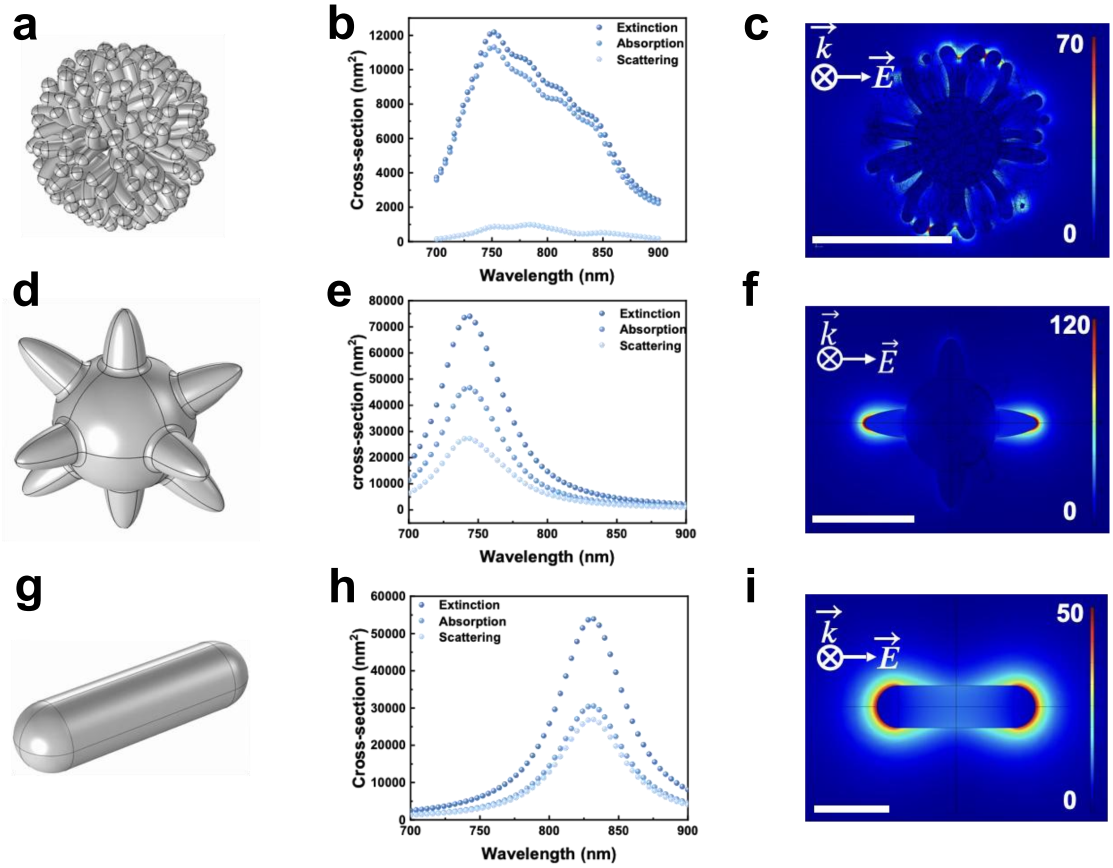


**Figure S7.** (a) The geometric model of HBGNCs (60 nm) for the theoretical simulation. (b) Scattering, absorption, and extinction cross-sections of HBGNCs. (c) Near-field distribution of HBGNCs at 800 nm. (d) The geometric model of AuNSTs (Core: 54 nm, branch: 20 nm) for the theoretical simulation. (e) Scattering, absorption, and extinction cross-sections of AuNSTs. (f) Near-field distribution of AuNSTs at 750 nm. (g) The geometric model of AuNRs (104*28 nm) for the theoretical simulation. (h) Scattering, absorption, and extinction cross-sections of AuNRs. (i) Near-field distribution of AuNRs at 835 nm. Scale bars, 50 nm. In all the simulation studies mentioned above, the calculation results are for a single gold particle dispersed in water.

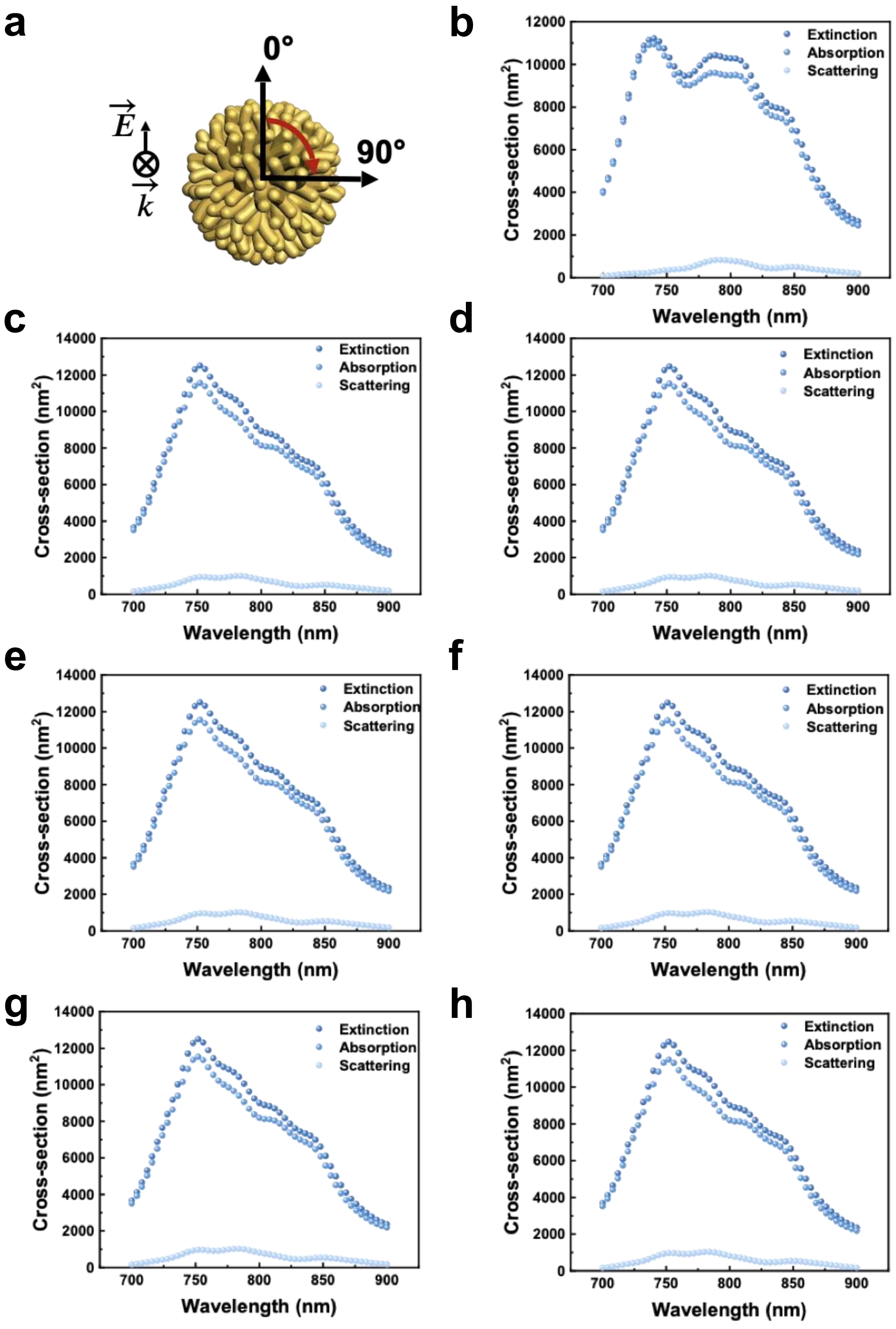


**Figure S8.** Calculated scattering, absorption, and extinction cross-sections of 62 nm HBGNCs at different polarization angles. (a) Schematic diagram of incident light. (b-h) The incident light angles are 0, 15, 30, 45, 60, 75, and 90 degrees, respectively. A single HBGNC was dispersed in water and excited by linearly polarized light at various polarization angles in the simulation studies.

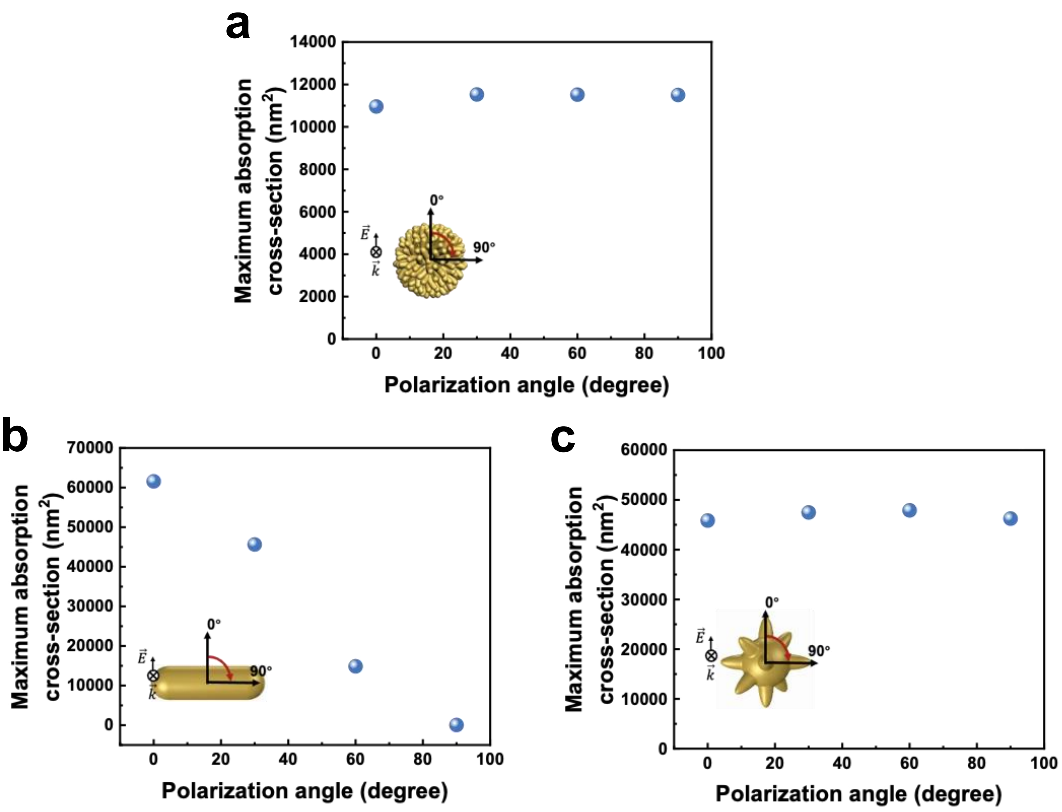


**Figure S9.** Maximum absorption cross-section at different light polarization directions for HBGNCs (a), AuNRs (b), and AuNSTs (c) within the 700-900 nm spectral range.

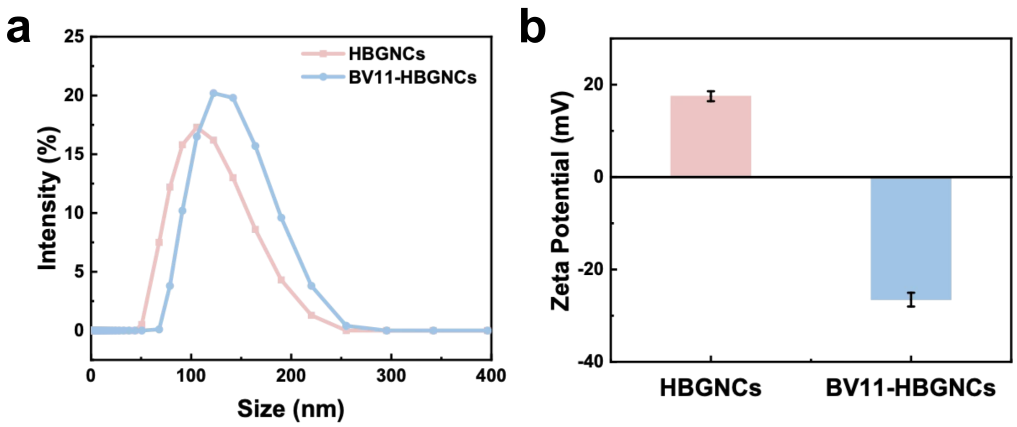


**Figure S10.** Characterization of mPEG-HBGNCs and BV11-HBGNCs using DLS. (a) Hydrodynamic diameters. (b) Zeta potentials.

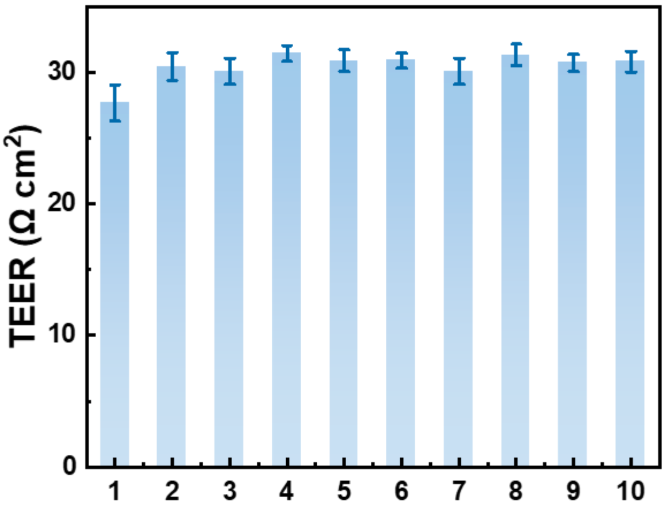


**Figure S11.** The transendothelial electrical resistance (TEER) of the in vitro BBB models (10 independent groups). Data are plotted as mean ± S.D., with 3 replicates in each experiment.

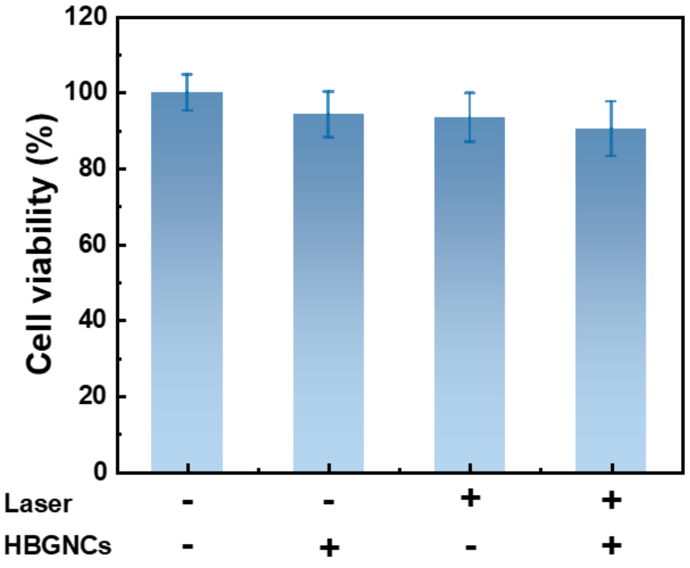


**Figure S12.** Cell viability under different experimental conditions. Data are plotted as mean ± S.D., with 3 replicates in each experiment. Concentration of the incubated HBGNCs solution: 0.04 nM. Laser excitation: 800 nm, 95 fs, 12.5 kHz, 60 μJ/cm².

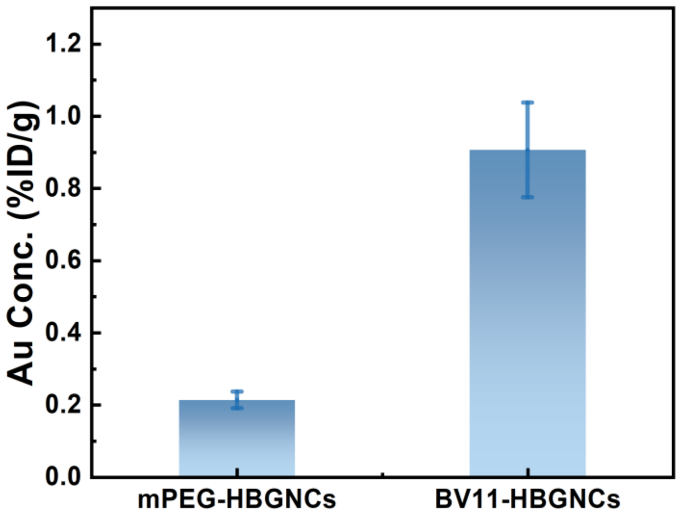


**Figure S13.** ICP-MS analysis of mPEG-HBGNCs and BV11-HBGNCs accumulation in the brain. Data are plotted as mean ± S.D., with 3 replicates in each experiment.

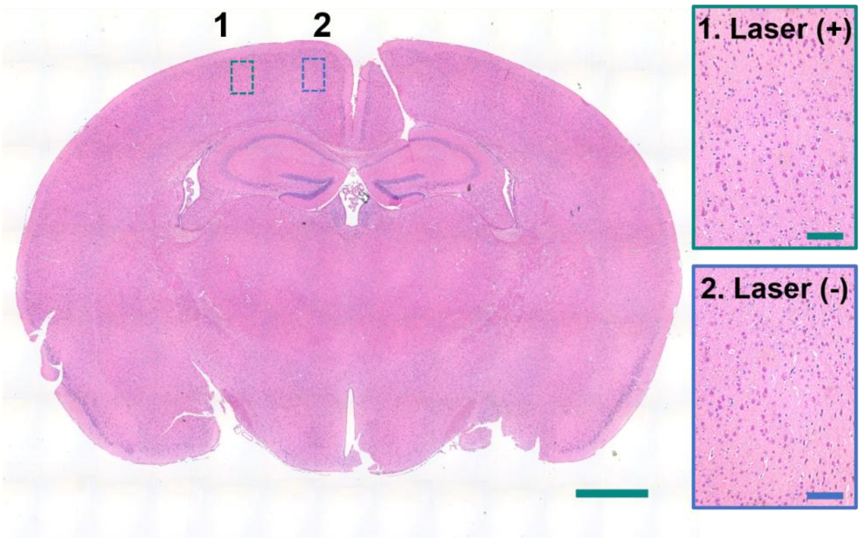


**Figure S14.** H&E staining of the brain tissue slices from the BV11-HBGNCs treatment group, with the enlarged views showing the irradiated cortex (1) and the contralateral nonirradiated cortex (2). Scale bars, 1 mm and 250 μm.

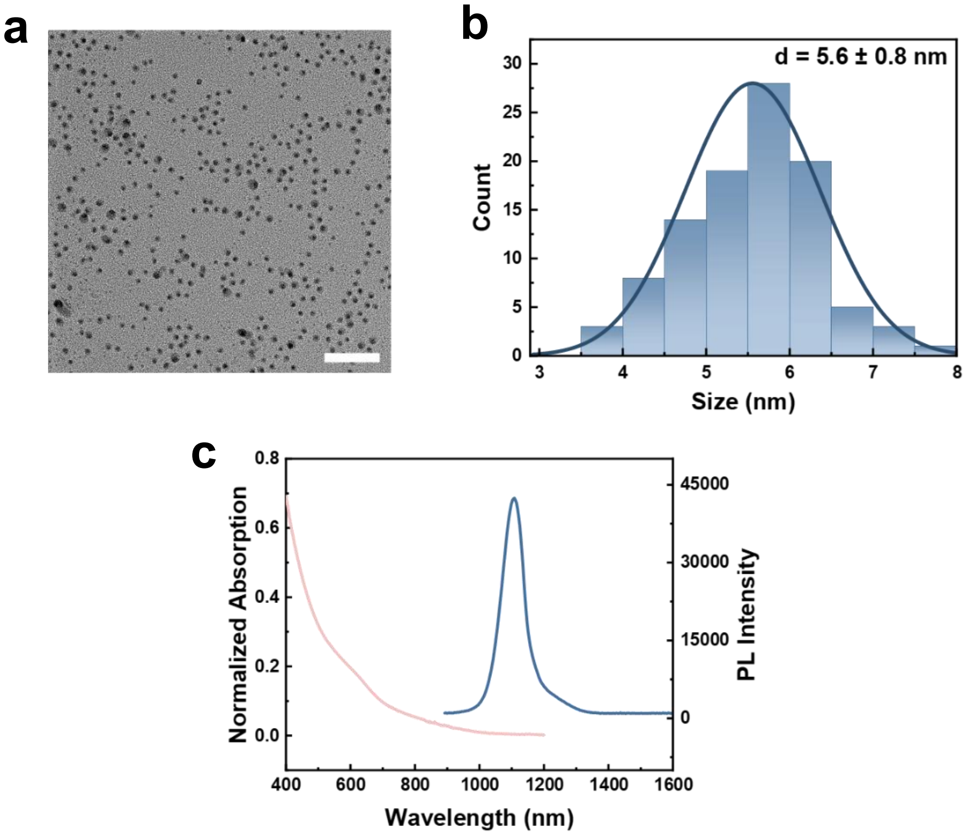


**Figure S15. (**a) TEM image of mPEG-AgAuSe QDs, scale bar: 50 nm. (b) size distribution histogram of mPEG-AgAuSe QDs. (c) Absorption and photoluminescence spectra of the mPEG-AgAuSe QDs.

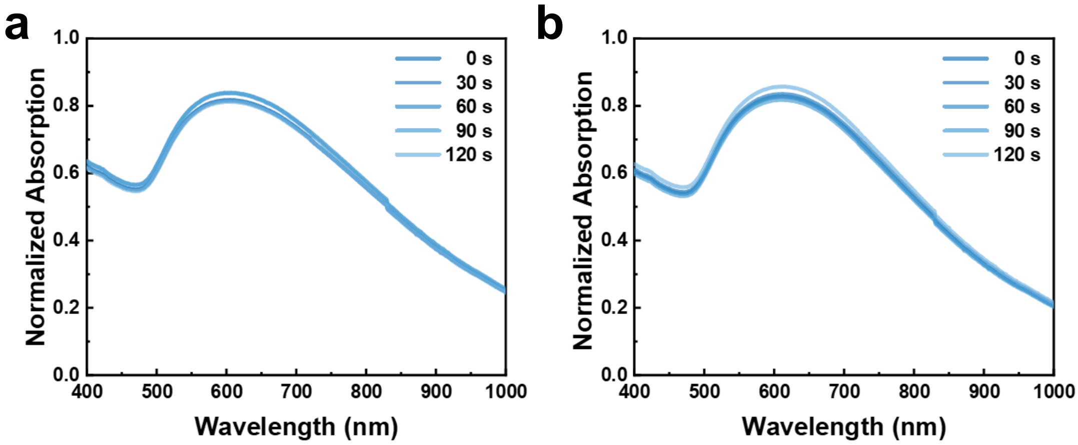


**Figure S16.** UV-vis-NIR spectra of 62 nm HBGNC under fs laser irradiation (800 nm, 12.5 kHz, 0.5 mJ/cm$^2$ (a) and 1.0 mJ/cm$^2$ (b)) for different durations.

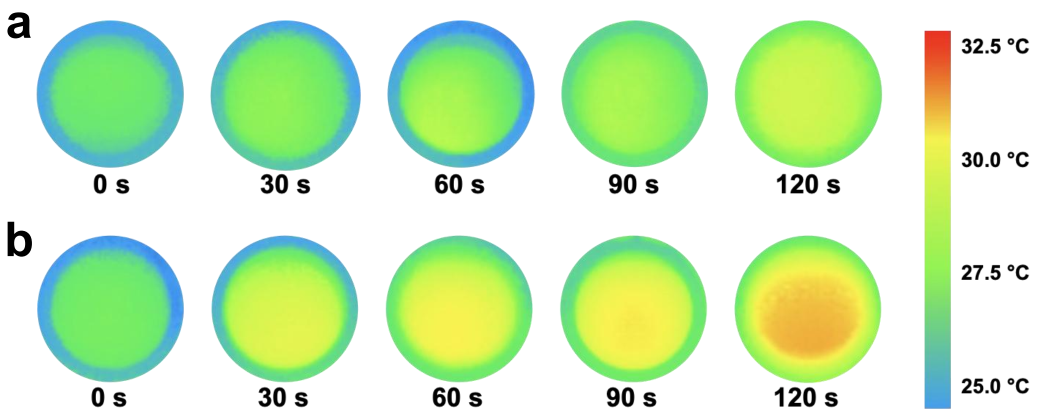


**Figure S17.** Thermal imaging maps of 62 nm HBGNC solution with absorbance of 3.5 at different time points under fs laser irradiation (800 nm, 12.5 kHz, 0.5 mJ/cm$^2$ (a) and 1.0 mJ/cm$^2$ (b)).

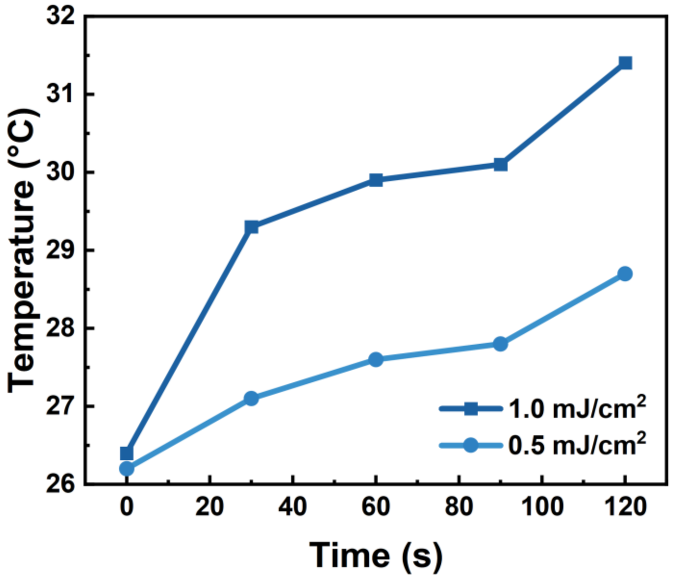


**Figure S18.** Temperature rise curves of 62 nm HBGNC solution with an absorbance of 3.5 after 120 seconds of 800 nm fs laser irradiation under different light intensities.
